# S-adenosylhomocysteine hydrolase regulates anterior patterning in *Dugesia japonica*

**DOI:** 10.1101/2020.01.22.916072

**Authors:** Kristina Reinmets, Johanna Bischof, Emily Taketa, Michael Levin, Stephen M. Fuchs

**Affiliations:** Department of Biology, Tufts University, Medford, MA, USA; Allen Discovery Center at Tufts, Tufts University, Medford, MA, USA; Institute for Protein Innovation, Boston, MA, USA

## Abstract

**Background:** Biological methylation requires S-adenosylmethionine (SAM) and participates in a range of processes from modulation of gene expression via histone modifications to neurotransmitter synthesis. An important factor in all methylation reactions is the concentration ratio of SAM to methylation byproduct S-adenosylhomocysteine (SAH). SAH hydrolase, also known as adenosylhomocysteinase, depletes SAH and thereby facilitates metabolite recycling and maintains the methylation permissive SAM/SAH ratio. While the importance of SAH hydrolase in sustaining methylation is obvious on the cellular level, the function of this metabolic process on the organismal scale is not clear.

**Results:** We used planarian *Dugesia japonica* to investigate the role SAH hydrolase in physiological homeostasis on the body-wide scale. Remarkably, pharmacological inhibition of the SAH hydrolase results in regression of anterior tissues and is accompanied by extensive apoptosis throughout the planarian body. Moreover, exposure to the SAHH inhibitor AdOx leads to changes in brain morphology and spatial shift in the expression of Wnt-modulator *Notum*. Strikingly, planarians are able to overcome these destructive patterning defects through regeneration of the anterior tissues and adaptation to the used inhibitor. Transcriptome analysis indicates that resistance to the SAHH inhibitor is at least partly mediated by changes in folate cycle and lipid metabolism.

**Conclusions:** SAH hydrolase plays a critical role in planarian homeostasis and anterior patterning potentially through modulation of Wnt signaling. Moreover, planarian adaptation to the SAHH inhibitor via metabolic reprogramming suggests potential targets for addressing methylation-related human conditions.

## Background

Biological methylation uses S-adenosylmethionine (SAM) and results in formation of a methylated product and S-adenosylhomocysteine (SAH). SAH is a potent methylation inhibitor and needs to be metabolized to sustain methylation-permissive conditions in the cells [1]. SAM-dependent reactions involve methylation of myriad substrates, from nucleic acids to proteins and lipids and thereby methylation regulates virtually every aspect of cell physiology [2]–[4]. Therefore, removal of SAH by SAH hydrolase (SAHH) sustains the balance of SAM to SAH ratio and provides a segue into metabolite recycling via homocysteine production (Fig. 1A) [5]. Homocysteine can be either recycled for SAM production or used in biosynthesis of the biological antioxidant glutathione. Perhaps unsurprisingly, changes in physiological levels of metabolites on either side of SAHH-mediated reactions are associated with human diseases. For example, elevated plasma SAH and homocysteine are associated with Alzheimer’s disease, dementia, cardiovascular as well as cerebrovascular diseases [6]–[10]. Moreover, importance of SAH hydrolase is reinforced by the fact that only four human cases have been reported of mutation in *SAHH* gene [11]–[14]. Vital function of this enzyme makes it difficult to study in complex organisms. Therefore, simple model organisms can bring to light much needed insight into the role of SAH hydrolase and related metabolites in health and disease.

**Figure 1.**
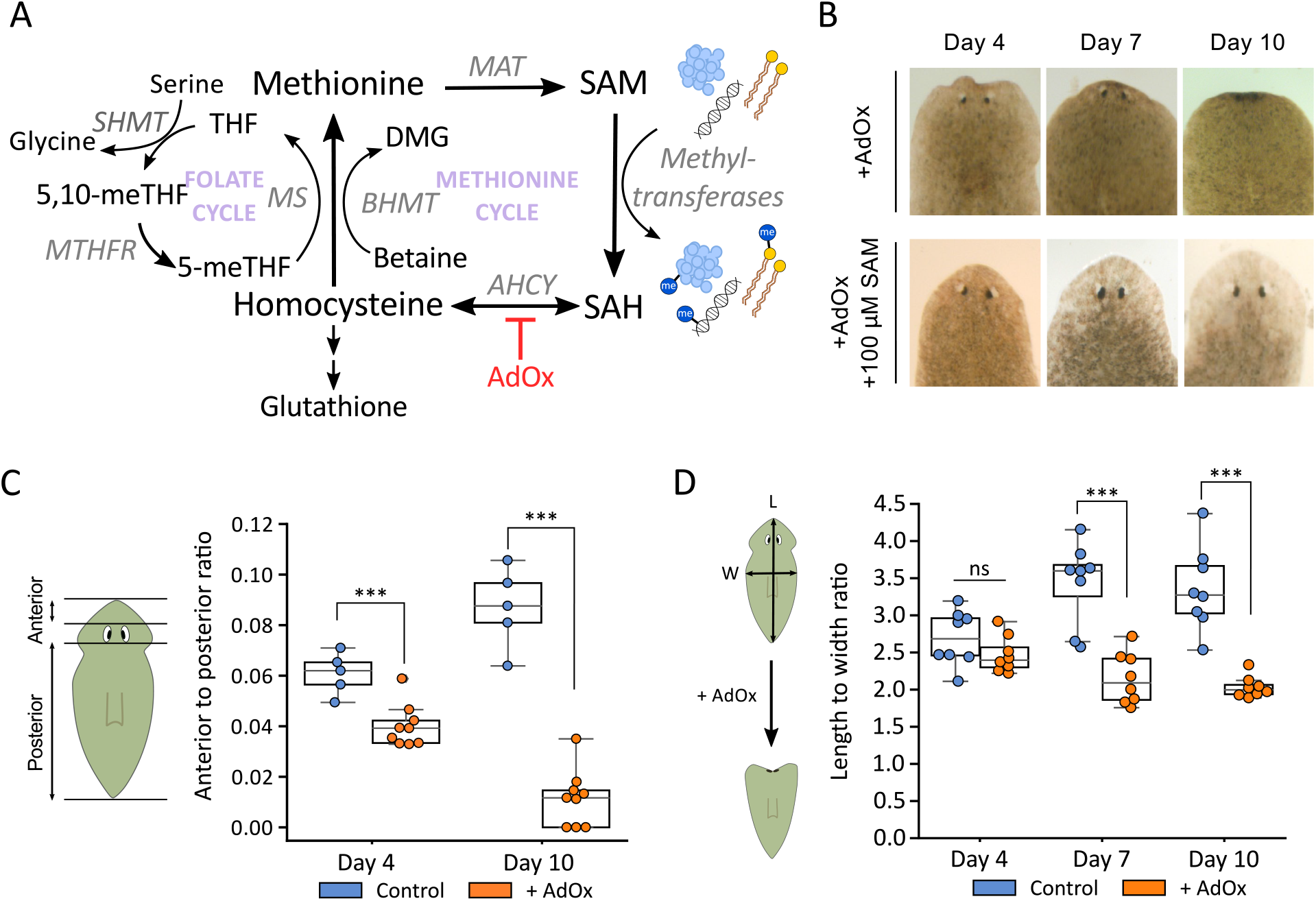
Inhibition of SAH hydrolase causes head regression and scaling defects in *D. japonica*. A – schematic of metabolic pathways producing and maintaining S-adenosylmethionine (SAM). Abbreviations: *MAT* – methionine adenosyltransferase, SAH – S-adenosylhomocysteine, *BHMT* – betaine homocysteine S-methyltransferase, DMG – dimethylglycine, THF – tetrahydrofolate, 5,10-meTHF – 5,10-methylenetetrahydrofolate, 5-meTHF – 5-methyltetrahydrofolate, *MS* – methionine synthase, *SHMT* – serine hydroxymethyltransferase, *MTHFR* – methylenetetrahydrofolate reductase. SAHH inhibitor AdOx shown in red. B – AdOx-induced head regression over time. Bottom row – phenotype is rescued by simultaneous supplementation with 100 μM SAM. C – anterior tissue loss shown as a ratio of distance anterlior versus posterior to the eye spots. D – changes in body length to width ratio during exposure to AdOx. *** p < 0.001, Student’s t-test.

Planarian flatworms have incredible metabolic flexibility exemplified by their ability to withstand starvation, regenerate missing tissues without food consumption, and adapt their body size based on nutrient availability. In fact, body length of planarians from the same species can differ up to 40-fold [15], [16]. Such metabolic robustness suggests that they could be a good model organism to look for treatment clues for metabolic diseases like the ones brought upon by SAHH-related metabolites. Despite the long-known metabolic power of planarian flatworms, metabolism of these organisms has only recently started gaining attention [16], [17].

In this work, we used planarian *Dugesia japonica* as a model to study the role SAH hydrolase in physiological homeostasis. We show that pharmacological inhibition of the SAH hydrolase results in head degeneration accompanied by body-wide apoptosis, changes in brain shape and spatial shift in expression of Wnt-antagonist *Notum*. However, planarians are able to overcome this severe phenotype by regenerating the anterior tissues and developing resistance to the SAHH inhibitor adenosine-dialdehyde. RNA-seq results show that resistance to the metabolic inhibitor is at least partly mediated by changes in genes from one-carbon and lipid metabolism. Further understanding of the mechanisms mediating the adaptation to the SAHH inhibition could provide potential targets for therapeutic interventions for methylation-related diseases.

## Results

### Inhibition of SAH hydrolase causes head regression in *D. japonica*

SAH hydrolase (SAHH) carries out a reversible hydrolysis of SAH – a critical reaction needed for sustaining the methylation-permissive conditions in cells. To investigate the importance this reaction in the planarian *D. japonica*, we used a specific and irreversible inhibitor of SAHH called adenosine-dialdehyde (AdOX) (Fig. 1A) [18], [19]. Inhibition of SAHH has multiple cellular consequences. First, it leads to accumulation of SAH which affects all methylation reactions via competition with SAM for the enzyme active site. Magnitude of this effect varies between different enzymes due to differences in substrate affinity [20]. Secondly, SAHH inhibition limits the production of homocysteine which is an important precursor for production of cysteine and glutathione. As methylation of myriad of substrates and redox maintenance are crucial for every cell, exposure of planarians to the SAHH inhibitor AdOx may be expected to affect the whole organism. Strikingly, intact planarians treated with 100 μM AdOX start showing a twitch-like movement of the head around day 4 of the drug exposure by pulling in the tip of the head (Fig. 1B). The twitch-like phenotype then progresses into head regression over the course of ten days leaving planarians with virtually no tissue anterior to the photoreceptors (Fig. 1B). Meanwhile, the remaining body does not show any obvious signs of tissue loss. AdOx-induced head regression can be rescued by co-administration of 100 μM SAM with the inhibitor, confirming that the observed phenotype is primarily caused by the inhibition of methylation reactions.

Loss of the anterior tissues in AdOx-treated planarians is readily confirmed upon fixation of the worms which causes muscle relaxation and eliminates the possibility of an inward contraction of the head as opposed to tissue loss. We measured the distance from the eyes to the tip of the head versus eyes to the tip of the tail (Fig. 1C). The resulting ratio of the anterior to posterior distances, respectively, shows significant reduction of the frontal tissues during AdOx exposure. Moreover, the anterior tissue loss during SAHH inhibition leads to changes in body proportions by decreasing the length to width ratio relative to control individuals (Fig. 1D). Planarians are known to be able to adjust their body size and remodel their organs based on nutrient availability as well as after fissioning (asexual reproduction) to maintain their set proportions [15]. Therefore, failure to maintain the correct proportions indicates significant perturbation to planarian physiology both in a region-specific manner in the head as well as throughout the body.

### AdOx-induced head regression is accompanied by ectopic pigment cup cell formation

The loss of anterior tissues is so dramatic it prompted us to focus on this region more closely. During our examination of the regressing region in AdOx-treated worms, we noticed the formation of dark pigments posterior to the photoreceptors. Coloration of those pigments resembled eye pigment cup cells. While three types of pigments – porphyrine, ommochrome, and melanin have been identified in planarians, melanin-based pigments are only produced by the pigmented optic cup cells of the planarian eye [21], [22]. These melanin pigments are the least sensitive to light exposure and remain detectable after standard bleaching procedures of exposure to light and hydrogen peroxide (Additional file: Fig. S1). The ectopic anterior pigments observed in AdOx-treated planarians remained visible after bleaching, suggesting that they are indeed melanin-based pigment cup cells. We wondered if these pigmented optic cup cells were part of ectopically forming eyes. To address this question, we used immunohistochemistry to look at planarian visual circuit neurons during head regression on day 7 (Fig. 2A). Curiously, the ectopic pigment cup cells were not accompanied by the photoreceptor neurons (PRN) which were only present in the eyes. This finding is unusual, as most reported supernumerary eyes are innervated [23], [24]. Moreover, the overall structure of the PRN-s did not seem to be affected by the drug treatment despite the ongoing head regression. Although we observed some pigment cup cells outside of the pigment cups in control individuals, the number of the ectopic pigment cup cells increased dramatically in AdOx-exposed planarians (Fig. 2B).

**Figure 2.**
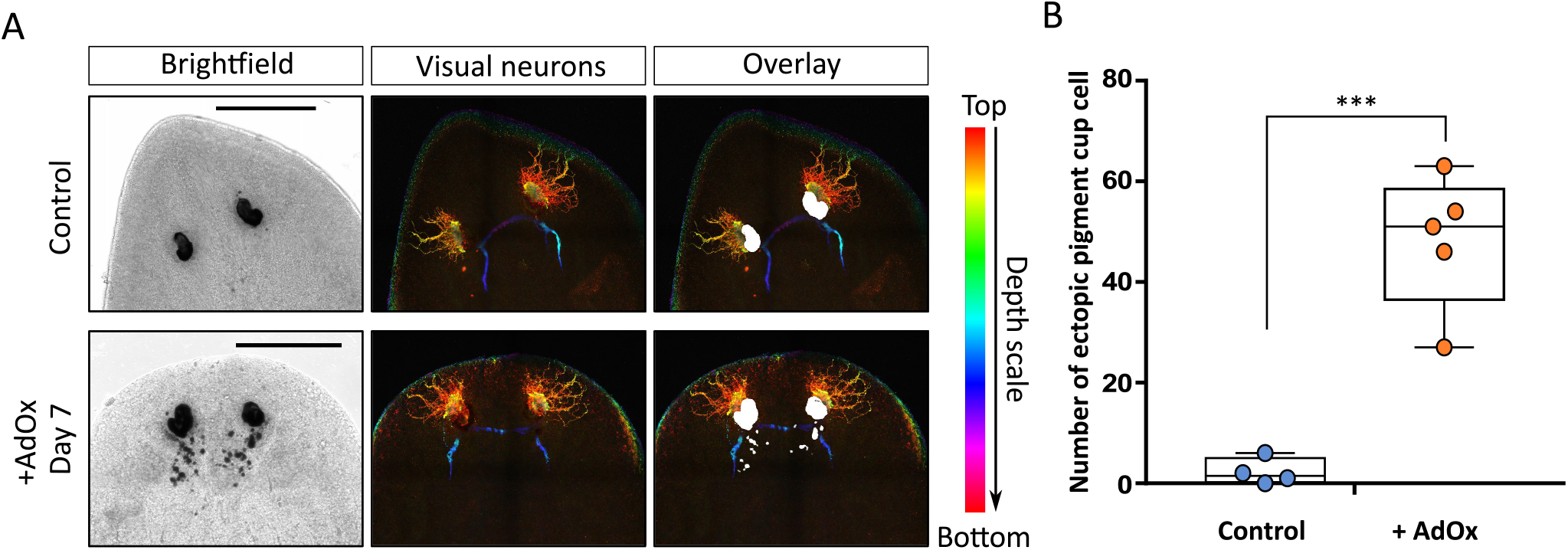
SAH hydrolase inhibition with AdOx causes formation of ectopic pigment cup cells. A – confocal images of visual circuit neurons visualized with anti-VC1 antibody in controls and planarians on Day 7 of the drug exposure. White structures on the overlay images are eye pigment cups. Scale bars – 200 μm. B – quantification of pigment cup cells outside of the pigment cup in controls and Day 7 AdOx-treated worms. *** p < 0.001, Student’s t-test.

### Inhibition of SAH hydrolase leads to systemic cell death and changes in brain morphology

We hypothesized that SAHH inhibition-induced head regression could be mediated by two different mechanisms – cell death or loss of proliferating cells. First, we explored the latter. In planarians, the only proliferating cells are adult pluripotent stem cells, called neoblasts, and their progeny [25]. Neoblasts reside in the mesenchymal space of the trunk and tail region but are absent from the head. Various manipulations that lead to defects in neoblast renewal are known to cause head regression phenotypes similar to the one we observed [26], [27]. Therefore, we used immunohistochemistry to visualize dividing cells as a proxy for neoblasts over the course of AdOx exposure using anti-H3P antibody (Additional file: Fig. S2). Interestingly, we saw no changes in localization or overall abundancy of the dividing cells, neither at the early stage of the phenotype on Day 4 nor on Day 10. Furthermore, the combination of AdOX with gamma-irradiation which specifically targets dividing cells did not intensify the head regression phenotype (Additional file: Fig. S3). Therefore, we conclude that SAHH inhibition causes head regression independently of neoblast proliferation defects.

Next, we explored the possibility of cell death-mediated head regression. Due to strong association of SAHH-related metabolites with neurodegenerative diseases, we decided to visualize apoptotic cells along with the central nervous system. To this end, we performed immunostaining using both an anti-caspase3 antibody to detect apoptosis and an anti-synapsin antibody for synaptic connections (Fig. 3A). Surprisingly, by the onset of the phenotype on day 4, six out of ten (6/10) immunostained individuals already showed increased cell death in the trunk region (Fig. 3A and C). Apoptosis signal was even stronger and more widespread on day 7 of the treatment when head regression was in progress but seemed to localize to the base of the brain by day 10. We observed similar results using TUNEL assay which detects late-stage apoptotic and necrotic cells undergoing DNA fragmentation (Additional file: Fig. S4). Strikingly, the AdOx-induced apoptosis was happening almost exclusively outside of the anterior-most tissue which was seemingly degenerating. Furthermore, even though integrity of the nervous system was not affected by this metabolic intervention in the early timepoints, morphology of the brain had changed by day 10 (Fig. 3A and B). Orientation of the characteristic side-branches on either lobe was nearly parallel relative to the body midline, and the brain appeared shorter and wider (compare Fig. 3B Control and 3B +AdOx, Day10). Quantification of the brain height to width ratio in control and AdOx-treated animals confirmed our observation of the changing brain shape (Fig. 3D). These results show that disruption of the methionine cycle via SAHH inhibition causes body-wide apoptosis and leads to morphological changes in the brain.

**Figure 3.**
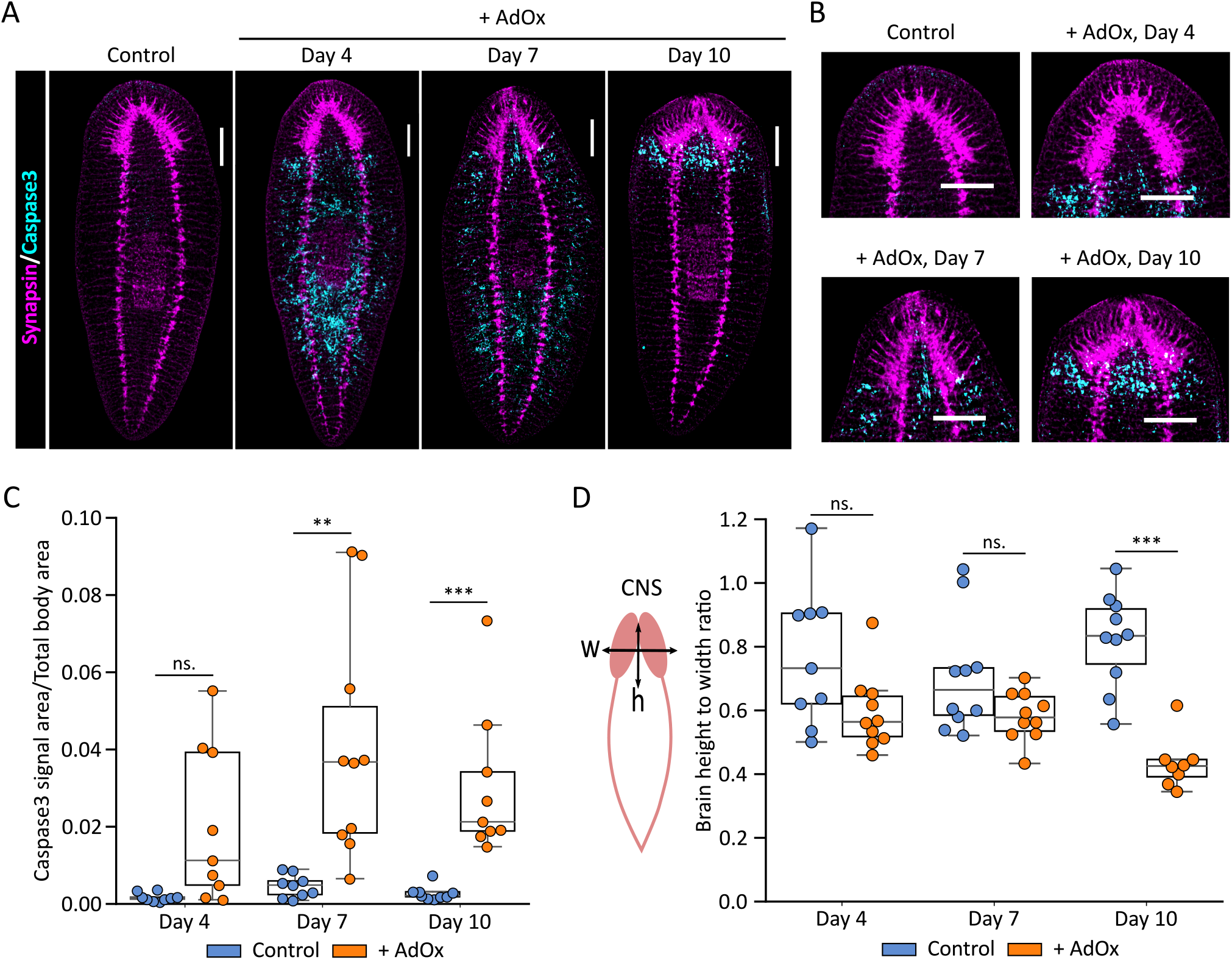
SAH hydrolase inhibition leads to body-wide cell death and changes in brain morphology. A – whole-mount double immunohistochemistry using anti-caspase3 and anti-synapsin antibodies visualizing apoptotic cells and planarian CNS, respectively. Caspase3 signal is shown in cyan and the nervous system is shown in magenta. Scale bars – 200 μm. B – zoomed view of images from A showing brain structure. C – caspase3 signal quantification showing ratio of caspase3-positive area over total body area. D – brain height to width ratio in control and AdOx-treated planarians. ** p < 0.01, *** p < 0.001, Student’s t-test.

### AdOx exposure affects spatial expression of anterior pole genes

The regressing head region in planarians treated with AdOx overlaps with the anterior pole – a region of cells expressing signaling molecules needed for regional specification. Expression of the anterior pole genes has an important role both in planarian regeneration as well as homeostatic tissue maintenance. We therefore hypothesized that SAHH inhibition may lead to head regression by affecting the required morphogen patterns in the head. We used fluorescent *in situ* hybridization to look at the expression of two genes – a Wnt-antagonist *notum* and FGFRL gene *ndl4* (Fig. 4A) – both well-established anterior markers in planarians [28]. Strikingly, *notum* was still expressed 10 days into AdOx exposure with the expressing cells being shifted posteriorly compared to control animals. *Notum* expression is normally limited to the anterior pole with some expression in the top of the brain, anterior to the eyes [29]. However, in AdOx-treated planarians, a dense population of *notum* expressing cells was present just above the eyespots. On the other hand, *ndl4*, which is normally expressed as a gradient from the tip of the head to the region posterior to the eyespots, seemed to be affected differently. We detected ndl4 expression in a narrower region compared to control individuals (Fig. 4A and B). These results show that SAHH inhibition changes the localization of *notum* expression but does not modulate the *ndl4* expression domain.

**Figure 4.**
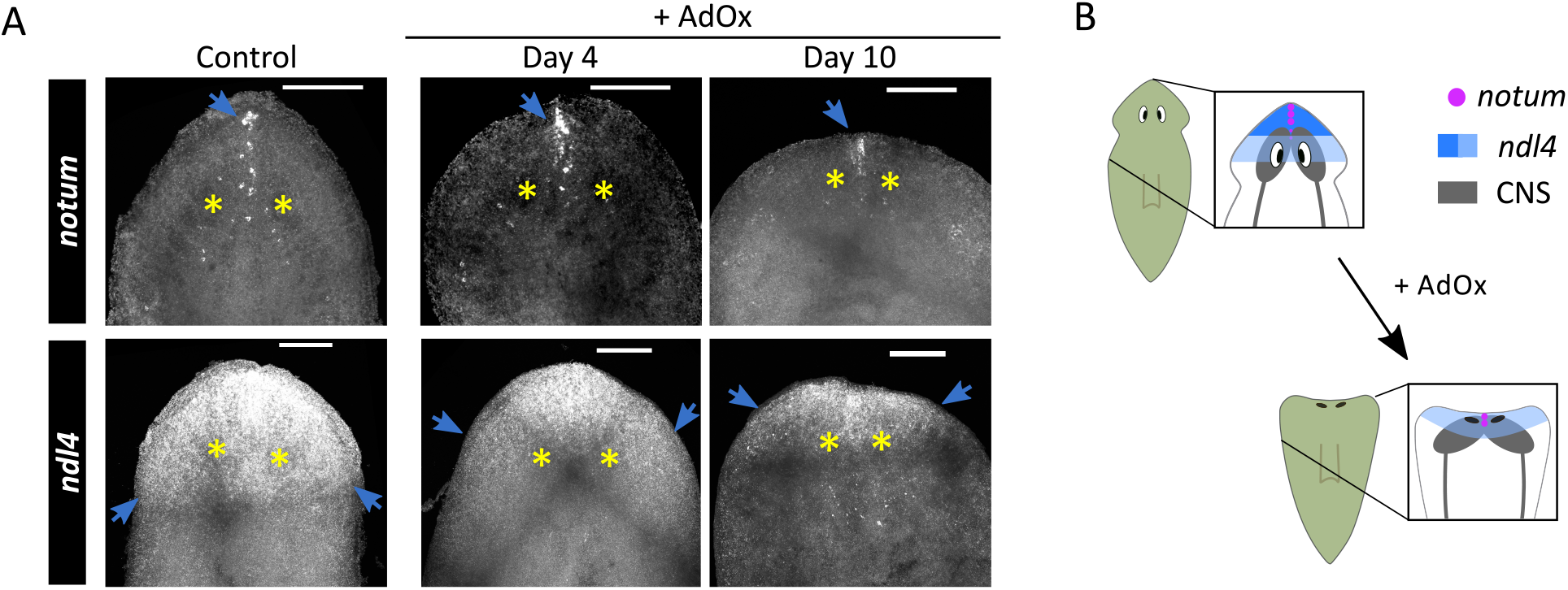
Spatial expression of anterior pole genes is disrupted by inhibition of SAH hydrolase. A – FISH-showing changes in spatial expression of *notum* and *ndl4* in control and AdOx treated planarians on days 4 and 10 of drug exposure. Yellow asterisks indicate eyespots, blue arrows point at the fluorescent signal. Scale bars – 200 μm. B – schematic summary of the FISH results showing how AdOx affects expression of anteriorly expressed genes.

### Planarians regenerate lost tissues despite SAHH inhibition

Given the robustness of planarian physiology in dealing with stressors like tissue loss, we continued to keep planarians in the original drug-containing solution without refreshing it past head regression. At around three weeks of AdOx exposure, we noticed the formation of anterior blastemas – a collection of undifferentiated cells which can give rise to new tissues, indicating regeneration (Additional file: Fig. S5). By one month, 83% of the planarians had restored normal head shape with 10% of the worms displaying abnormal growth, while 7% did not regenerate (Fig. 5A and B). Interestingly, some planarians had retained the old eye pigments in addition to the newly regenerated eyes. We performed immunohistochemistry to assess whether those original pigment cups still contained photoreceptor neurons (Fig. 5C). We found that new and not the original pigment cups contained the photoreceptor neurons and thus only the new eyes are likely to be functional. Remarkably, number of ectopic pigment cup cells observed during the head regression on Day 7 had declined in some but not all (3 out of 5) examined individuals (Additional file: Fig. S6). Moreover, the regenerated planarians also seemed to have overcome the body scaling defects observed during the head regression and resembled control worms in their length to width proportions (Fig. 5D). Finally, having observed severe changes in the brain during AdOx-induced head regression, we wondered if planarians had been able to overcome the neurotoxicity of SAHH inhibition. We visualized synaptic connections by immunohistochemistry to look at the CNS and observed a range of phenotypes from healthy looking brains to wide diffused lobes (Fig. 5E). One out of ten assayed individuals displayed abnormal secondary cephalic connection (Fig. 5E +AdOx, rightmost image). However, the shape of the brains from AdOx-treated worms had improved compared to Day 10 of the drug exposure based on the height to width ratio (Additional file: Fig. S7).

**Figure 5.**
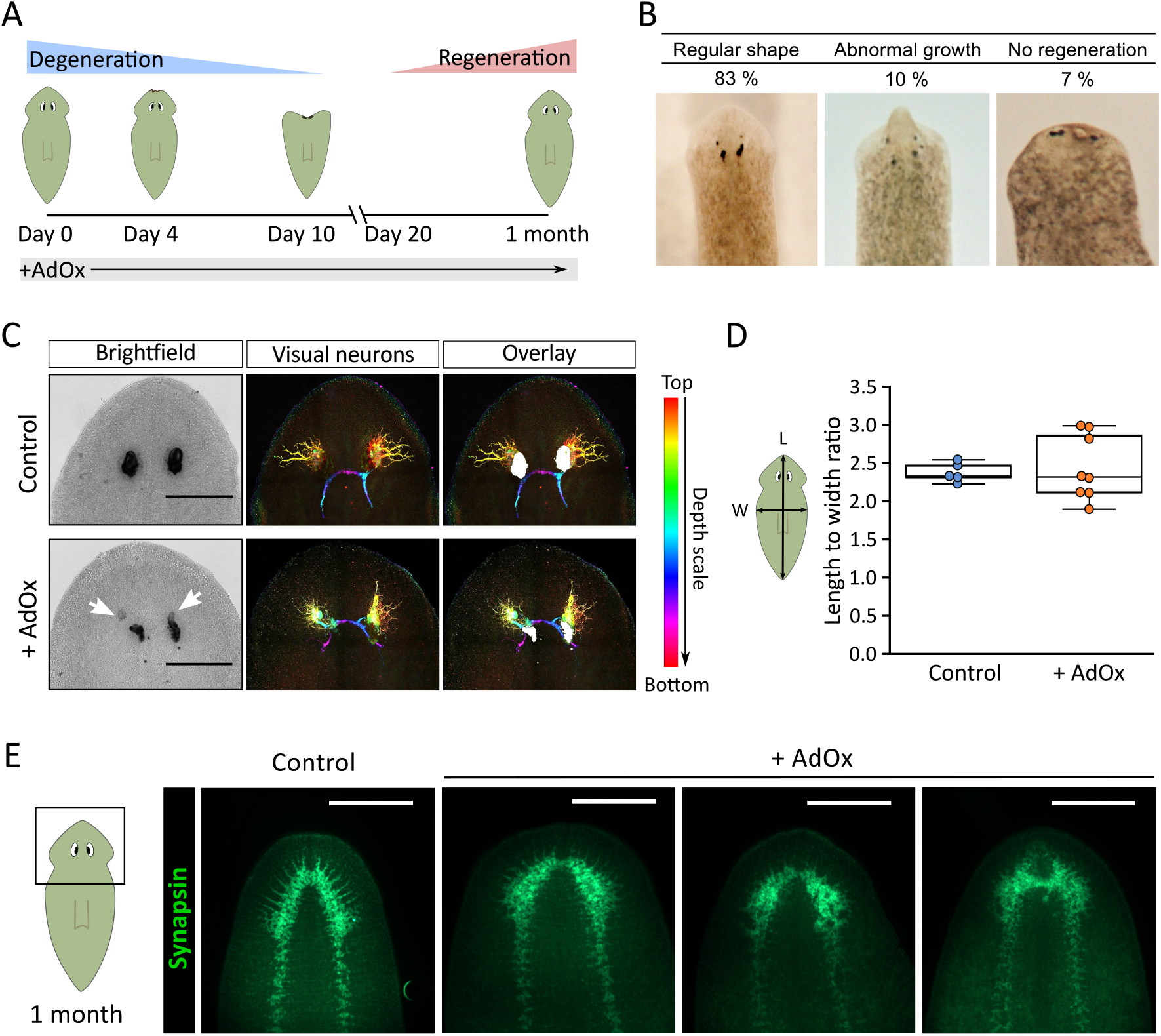
Planarians restore anterior tissues despite the SAHH inhibition. A –schematic and timeline of changes planarians undergo during the exposure to AdOx. B – head phenotypes of planarians after 1 month of AdOx exposure. C – visual neurons of AdOx-exposed worms at 1 month timepoint. Visual neurons visualized with anti-VC1 antibody. White arrows on brightfield images indicate the regenerating pigment cups. White structures on shown on the overlay images are eye pigment cups. D – length to width ratio of control and 1 month AdOx individuals. E – immunostaining of the CNS using anti-synapsin antibody. Scale bars – 200 μm.

### Planarians are resistant to AdOx upon second exposure

Surprised by the recovery of AdOx-exposed planarians, we were curious whether the regenerated individuals would have altered sensitivity to the inhibitor if subjected to another round of treatment. Unexpectedly, we saw no head regression when planarian water was refreshed with the same concentration of the inhibitor after one month (Fig. 6A). We have previously observed a similar adaptation of *D. japonica* to BaCl_2_ which induces very rapid (within 3 days) head degeneration while AdOx-prompted head regression happens relatively slowly (within 10 days) [30]. We first hypothesized that adaptation to AdOx arises due to exposure to the inhibitor during head regeneration. We performed a head amputation experiment and kept the resulting fragments in the same concentration of AdOx as during the long-term experiments. Planarians were able to regenerate both anterior and posterior tissues despite the SAHH inhibition (Additional file: Fig. S8). However, after the initial head regeneration was completed, planarians exhibited the same AdOx-induced head regression as the intact individuals. This suggests that adaptation to AdOx occurs following the drug-induced head regression.

**Figure 6.**
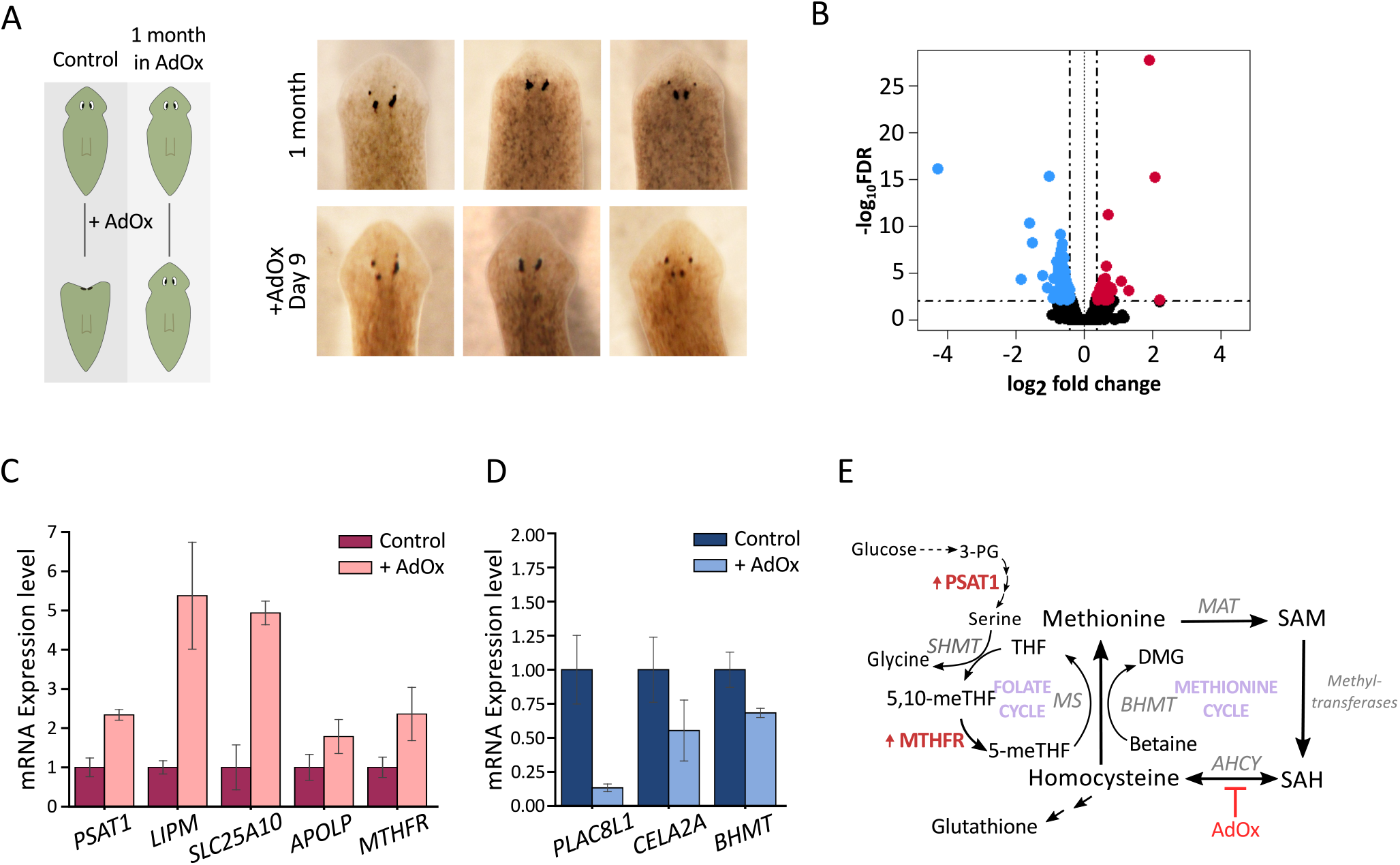
Planarians are resistant to AdOx upon second exposure. A – prolonged exposure to AdOx leads to acquired adaptation. B – volcano plot of RNA-seq-identified differentially expressed genes in AdOx-adapted planarians relative to control worms. Horizontal dashed line – adjusted p-value = 0.01, vertical dashed lines – log2 of −0.4 or 0.4. C – confirmation of top upregulated genes by qPCR (n = 2, error bars – std. error). D – confirmation of selected downregulated genes by qPCR (n = 2, error bars – std. error). E – schematic representation of one-carbon metabolism with genes upregulated in AdOx-resistance shown in red and indicated with “up”-arrows. *PSAT1* – phosphoserine aminotransferase 1.

### Adaptation to the SAHH inhibitor is mediated by the upregulation of one-carbon and lipid metabolism genes

To unravel the mechanism of this striking drug resistance, we performed an RNA-seq experiment on planarians kept in AdOx solution for 1 month and on respective controls. We found 119 significantly changing transcripts (FDR=0.01, log_2_=0.4 threshold, Fig. 6B, Additional file: Fig. S9, Table S1). Importantly, we found changes in key enzymes of one-carbon metabolism among the top differentially expressed genes (Fig. 6E). Specifically, we identified upregulation of *MTHFR* from the folate cycle and *PSAT1* which is involved in serine biosynthesis and can provide methyl-groups for the folate cycle (Fig. 6C). We also found upregulation of genes involved in lipid metabolism, *LIPM* and *APOLP*, as well as mitochondrial dicarboxylate transporter *SLC25A10*. Meanwhile, expression of *BHMT*, which participates in homocysteine re-methylation in the methionine cycle, was downregulated in AdOx-adapted planarians (Fig. 6D). We then set to confirm the sequencing results in an independent experiment. We were able to confirm the selected set of genes identified as upregulated by qPCR, although the expression level had changed by different magnitude (Additional file: Fig. S10). This shows that adaptation to the SAHH inhibition is consistently linked to upregulation of one-carbon and lipid metabolism. Given that organismal metabolism is highly affected by the availability of nutrients and AdOx treated planarians were starved, we hypothesized that metabolic reprogramming in AdOx-adapted worms may be affected by feeding. A group of planarians exposed to the SAHH inhibitor for 1 month was fed with liver paste twice and then placed into fresh drug solution. Strikingly, around 30 percent of the fed planarians (16/53) had become re-sensitized to the inhibitor, while all continuously starved individuals remained resistant to AdOx.

During our attempt to independently confirm a set of downregulated genes, we were able to verify only one out of four tested genes found as downregulated – a viral transcript with the highest sequence homology to *Solenopsis invicta virus-1* (SINV-1) (Additional file: Fig. S11). Interestingly, we found the same viral transcript mapping to two different contigs among our top up- and downregulated genes (Additional file: Table S1). Upon closer examination of the two contigs and planarian database (PlanMine [31]) entry they map to, it appeared that one of the two sequences was 4000 nucleotides longer. The longer transcript was the top downregulated gene in AdOx-adapted worms with almost 20-fold decrease in expression. Meanwhile, the shorter transcript was second most upregulated with 4-fold higher expression compared to controls. Notably, we were able to confirm this differential expression of the two SINV-1 transcript variants by qPCR both in the original sequencing experiment and in the independent replicate (Additional file: Fig. S11 and Fig. S12). Such drastic decrease in the longer transcript variant, *SINV-1L*, in AdOx-adapted planarians may suggests potential role for methylation in planarian susceptibility to viral infections.

## Discussion

In this study, we showed that SAH hydrolase has a critical role in planarian body homeostasis and its pharmacological inhibition leads to regression of the anterior tissues. This drastic phenotype appears to be mediated by body-wide apoptosis and leads to changes in brain morphology. Moreover, exposure to the SAHH inhibitor AdOx leads to spatial shift in expression of secreted Wnt antagonist *Notum*, moving it closer to the eyespots compared to control individuals. Remarkably, planarians become adapted to the SAH hydrolase inhibitor and resistant worms show upregulation of genes related to one-carbon and lipid metabolism.

In planarians, head regression is often associated with loss of stem cells or neoblasts, the only dividing cells in the body [25]. In this work, we did not observe significant changes in the number of dividing cells in AdOx-induced head regression. Although, there appears to be a slight decrease in the number of mitotic cells on day 10 of AdOx exposure. Interestingly, it has been shown that SAH hydrolase is required for proliferation of mouse embryonic stem cells [32]. This discrepancy could be either due to species-specific differences in stem cell metabolic demands or due to limitations of the pharmacological approach. However, silencing of the metabolic genes via RNAi is problematic due to their abundant expression. Deletion or mutation of the *Dj-SAHH* gene would be a more robust approach, however, molecular tools for generation of transgenic planarians do not exist yet. Furthermore, human and mouse embryonic stem cells have differential dependence on methionine versus threonine for 1-carbon units, respectively [33], [34]. Therefore, determining the metabolic demands specific to planarian stem cells will be necessary for better translation the current findings.

Our observation of the extensive apoptosis throughout the planarian body during AdOx-exposure exemplifies one of the strengths of planarians as a model system – their small size allows to monitor the molecular changes throughout the organism. Despite the apparent sensitivity of the anterior-most region of the worm, the apoptosis signal is not detected in that region. Instead, SAHH inhibition leads to systemic and not head-specific cell death. However, it is possible that rapid cell death in the body causes apparent deformation of the anterior-most tissues via loss of the supporting cell mass. On the other hand, formation of the ectopic pigment cup cells suggests potential disruption of the body plan maintenance. Moreover, spatial shift in the expression of secreted Wnt-antagonist *Notum* raises interesting questions. Are *Notum*-expressing cells migrating posteriorly or is *Notum* induced in this new more posterior location? It is known that *Notum* is regulated by β-catenin and change in spatial expression of *Notum* suggests potential changes in β-catenin activity [35]. Given the recently shown importance of methylation and one-carbon metabolism in canonical Wnt signaling by De Robertis group, we speculate that disruption of the methionine cycle may lead to failure in planarian body plan maintenance via changes in Wnt signaling [36], [37]. Although we did not find β-catenin target genes among differentially expressed genes in AdOx-adapted worms, it does not exclude possible changes in post-translational regulation. Further research is needed to elucidate the role of the SAHH and related metabolism in body-wide regulation of Wnt signaling.

Interesting questions are instigated by the planarian regenerative response to AdOx without exogenous damage. What initiates the head regeneration and why does it happen only after severe changes to anterior morphology? As we show by caspase3 staining, AdOx-induced apoptosis takes place outside of the planarian head region. Therefore, there is virtually no anterior tissue loss that could trigger head regeneration. Yet, planarians are able detect and fix these changes to their body plan. Understanding the mechanism behind such internal tracking of the shape and form will reveal potential ways of activating regeneration and inducing healing in other organisms.

Finally, planarian adaptation to AdOx provides valuable insight into possible modulation of metabolism in methylation-associated diseases. Upregulation of *MTHFR* and *PSAT1* genes provides a more efficient way to produce and transfer methyl-groups needed for SAM synthesis. While SAHH deficiency is extremely rare (four reported cases), aberrant levels of the substrate – SAH, as well as the product – homocysteine, of this enzyme strongly correlate with neurodegenerative and cardiovascular diseases. Therefore, changes in *MTHFR* and *PSAT1* genes in AdOx-adapted planarians suggest that perhaps stimulating the folate cycle could be beneficial in overcoming SAHH-related toxicity. Moreover, AdOx-adapted worms showed upregulation of *SLC25A10* – a mitochondrial dicarboxylate carrier which is reportedly needed for mitochondrial GSH transport in neuronal cells [38], [39]. Although the exact mechanism of GSH transport by SLC25A10 is unclear, it suggests that AdOx-treated worms may experience oxidative stress. Interestingly, *PSAT1* and *SLC25A10* are overexpressed in various cancer types where they promote cell proliferation and correlate with poor prognosis [40]–[43]. Hence, upregulation of these genes in AdOx-exposed planarians may help promote cell proliferation to counteract the extensive apoptosis triggered by the inhibitor. Overall, our results demonstrate the incredible plasticity of the planarian physiology in dealing with metabolic stressors.

## Conclusions

Taken together, we show that SAH hydrolase activity is critical for the homeostasis and body plan maintenance of the planarian *D. japonica*. Furthermore, the robust physiology of the planarians allows them to adapt to the SAH hydrolase inhibitor via upregulation of one-carbon and lipid metabolism-related genes, suggesting potential avenues for addressing the SAHH-associated human conditions.

## Materials and methods

### Planarian husbandry

An asexual strain of *Dugesia japonica* was maintained at 20 °C in the dark in Poland Spring® water and fed with organic calf liver paste once a week as described in [44]. All planarians used in this work were starved for at least 1 week before the start and during the experiments.

### Treatment with AdOx and SAM supplementation

Adenosine dialdehyde (AdOx) was obtained in solid form from Sigma-Aldrich (A7154) or Cayman Chemical (#15644). Fresh liquid stock of 90 mM concentration was prepared as per manufacturers’ instructions in 0.14 M HCl prior to every experiment and diluted in Poland Spring® to a final concentration of 100 μM. Planarians were kept in AdOx-containing or same dilution of HCl water throughout the experiments. For S-adenosylmethionine (SAM) supplementation, planarians were kept in Poland Spring water with 100 μM AdOx and 100 μM SAM (US Biological), 100 μM SAM only or control water.

### Irradiation

Animals were irradiated using Nordion Gamma Cell 1000 Irradiator with a Cesium-137 radiation source with a dose of 20 Gy achieved via 30 min exposure. Irradiated worms were then placed into 100 μM AdOx containing or control water and kept at 20 °C as in other experiments.

### Whole-mount immunohistochemistry

Planarians were fixed with Carnoy’s fixative (ethanol, chloroform, and glacial acetic acid at 6:3:1 ratio), bleached and stored in methanol as described in [45]. Fixed animals were gradually rehydrated into PBSTx.3 (PBS + 0.3% Triton X-100, pH 7.4) from 100% methanol by 25% increment and blocked in PBSTx.3BG (PBSTx.3 + 0.3% BSA + 10% goat serum) overnight at 4 °C. Blocking solution was then replaced with primary antibody diluted in PBSTx.3BG, incubated overnight at 4 °C, washed 6 times for 30 min with PBSTx.3B at room temperature and incubated with secondary antibody overnight at 4 °C. Samples were then rinsed 6 times with PBSTx.3B at room temperature and mounted in Vectashield® (Vector Laboratories) for subsequent imaging. The following primary antibodies were used in this study – rabbit-anti-H3p (MilliporeSigma, Cat # 04-817-MI) at 1:250 dilution, mouse-anti-synapsin (SYNORF1) 3C11 at 1:50 dilution (DSHB [46]), rabbit-anti-caspase-3 (Abcam, ab13847) at 1:300 dilution, and anti-Dj-arrestin (VC-1) at 1:1000 dilutionv[47]. Secondary antibodies were goat anti-mouse Alexa Fluor®594 (Invitrogen, Cat #A-11005) 1:400 dilution for synapsin, goat anti-rabbit Alexa Fluor®488 (Invitrogen, Cat #A-11008) at 1:400 dilution for caspase3, and goat anti-rabbit-HRP (Invitrogen, Cat # 65-6120) at 1:750 amplified with Alexa Fluor™ 555 Tyramide SuperBoost™ kit (Invitrogen, Cat # B40923) for H3p.

### Fluorescent *in situ* hybridization

Fluorescent riboprobes were synthesized as previously described [48]. Formaldehyde fixation and *in situ* hybridization was done as described in [49] with minor modifications. Namely, planarians were bleached for 1 hour using formamide as described in [48].

### Imaging and image analysis

Live planarians and H3p immunostained samples were imaged on Nikon SMZ1500 stereoscopic microscope (Nikon) with a Retiga 2000R camera (Teledyne QImaging) and Q-Capture software. Caspase3 and synapsin immunostained samples were imaged on Olympus BX61 (Olympus) using Hamamatsu ORCA AG CCD camera and Metamorph software. VC-1 immunohistochemsitry and FISH samples were imaged on a Leica SP8 confocal microscope (Leica Microsystems).

All image analysis was performed using FiJi. Briefly, quantification of H3p and caspase3 signal was done by setting the threshold and using Analyze Particles function. Caspase3 and synapsin images were merged using Merge Channels function. Post-processing of the confocal images was done by adjusting the contrast and interpolation of Z stacks with Z project function. Brain height to width measurements were done on the anti-synapsin immunohistochemistry images. Brain height was measured as distance between the anterior connection of the two lobes and the base of the brain. Brain width was measured as distance between anterior-most edges (or the 6^th^ branches) of the clustered branches [50].

### RNA-seq and transcriptome analysis

Planarian total RNA was isolated using TRIzol®Reagent (Invitrogen, Cat # 15596018) and 1-bromo-3-chloropropane (TCI America, Product # B0575), precipitated with isopropanol and treated with DNase I (Invitrogen, TURBO DNA-*free*™ Kit, Cat # AM1907) to remove DNA contamination. Sequencing library was prepared using NEBNext® Ultra™ II RNA Library Prep Kit for Illumina® (New England Biolabs Inc., Cat # E7770S) following manufacturer’s protocol. DNA concentration of the resulting libraries was measured using Qubit® 2.0 Fluorometer and Qubit™ 1x dsDNA HS Assay Kit (Invitrogen, Cat # Q33230) and all samples were analyzed on Fragment Analyzer (Agilent Technologies, Inc.).

Sequencing was done as Single End 150 on Illumina NextSeq550 instrument at Tufts University Genomics core facility. RNA-seq reads were mapped to *Dugesia japonica* transcriptome [51] using bowtie2 package and differential expression analysis was conducted with DESeq2 and Bonferroni-Holm adjusted p-value of 0.01 was used as a threshold to select significantly changing transcripts [52], [53].

### Quantitative RT-PCR

Total RNA was extracted using TRIzol®Reagent as described above and cDNA was prepared using SuperScript® First-Strand kit (Invitrogen, Cat # 11904018). QPCR was performed on ABI 7300 instrument (Applied Biosystems) using Brilliant III Ultra-Fast SYBR® QPCR Master Mix and custom primers (Additional file: Table S2).

## Contributions

K.R., J.B., and E.T. conducted the experiments. All authors conceived and designed the experiments as well as wrote and edited the manuscript.

## Acknowledgements

We would like to thank Dr. Devon Davidian for providing the riboprobe for Dj-*notum* gene used in the FISH experiment.

We are grateful to the Paul G. Allen Frontiers Group award (# 12171) and the NIH Research Infrastructure grant (NIH S10 OD021624) for supporting this research and members of the Allen Discovery Center at Tufts for general advice and technical support.

