## Supplementary figures and images for "S-adenosylhomocysteine hydrolase regulates anterior patterning in *Dugesia japonica*"

### Fig. S1

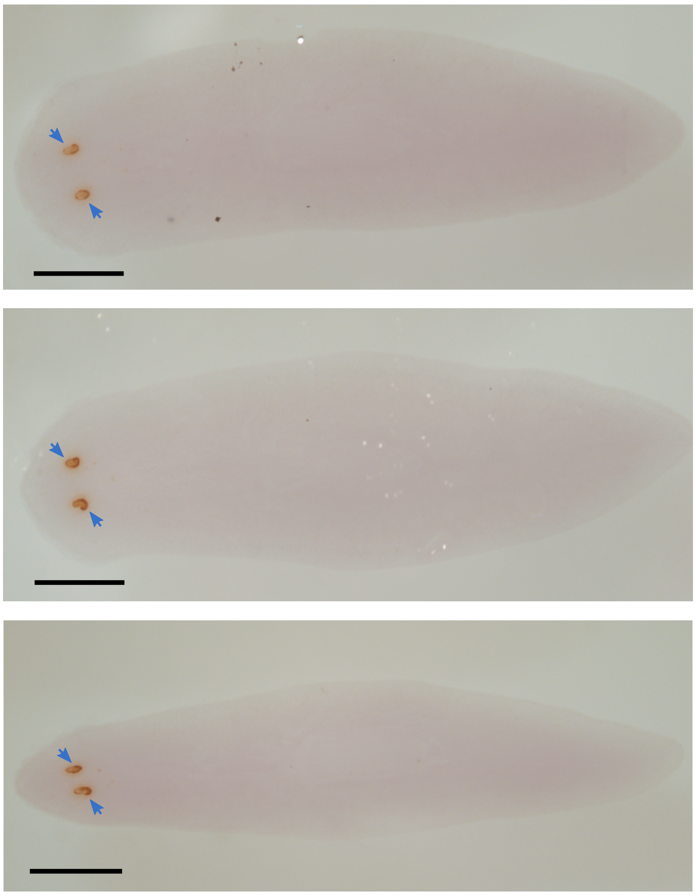

**Figure S1.**

### Fig. S2

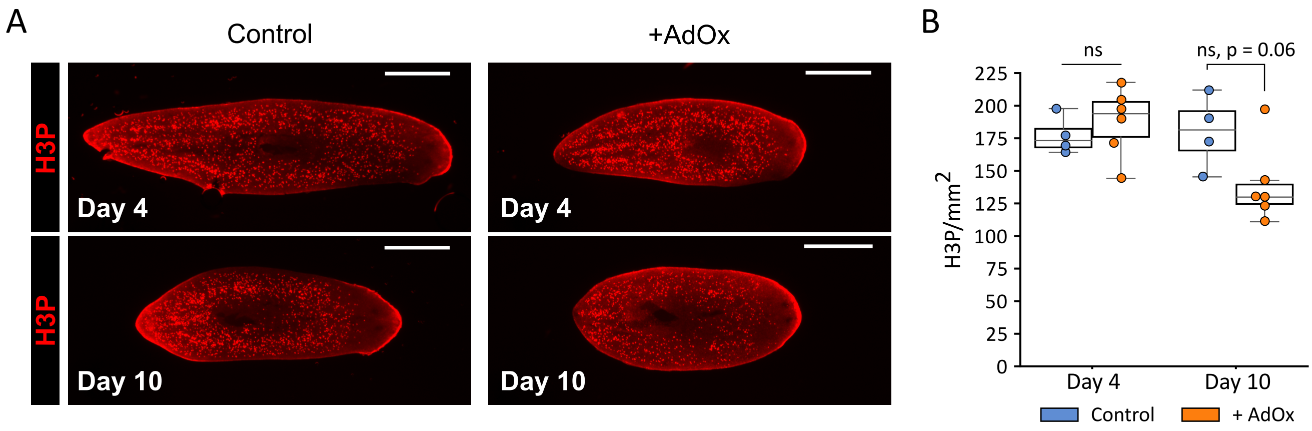

### Fig. S3

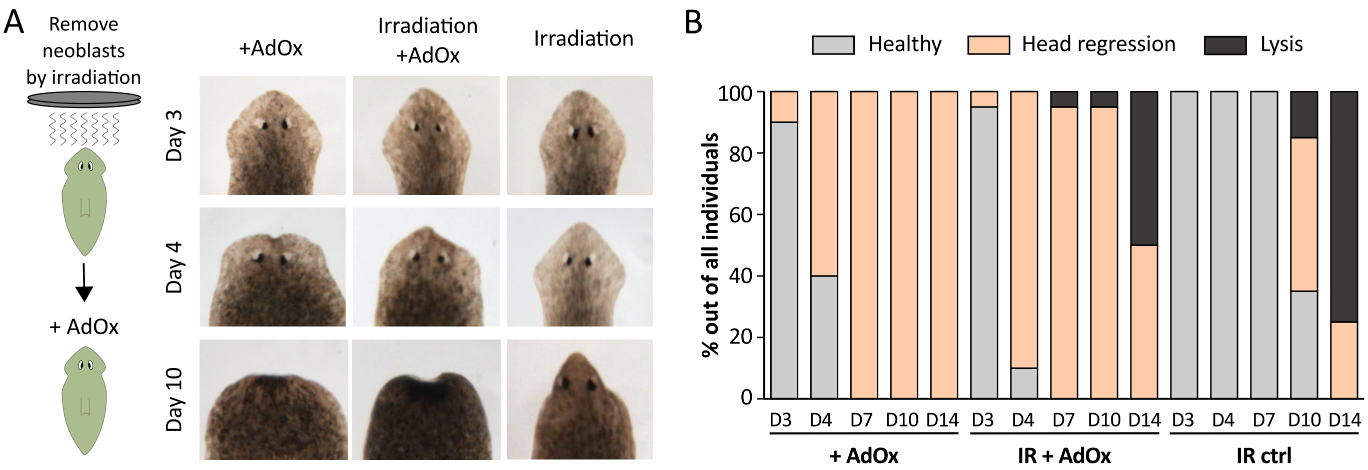

### Fig. S4

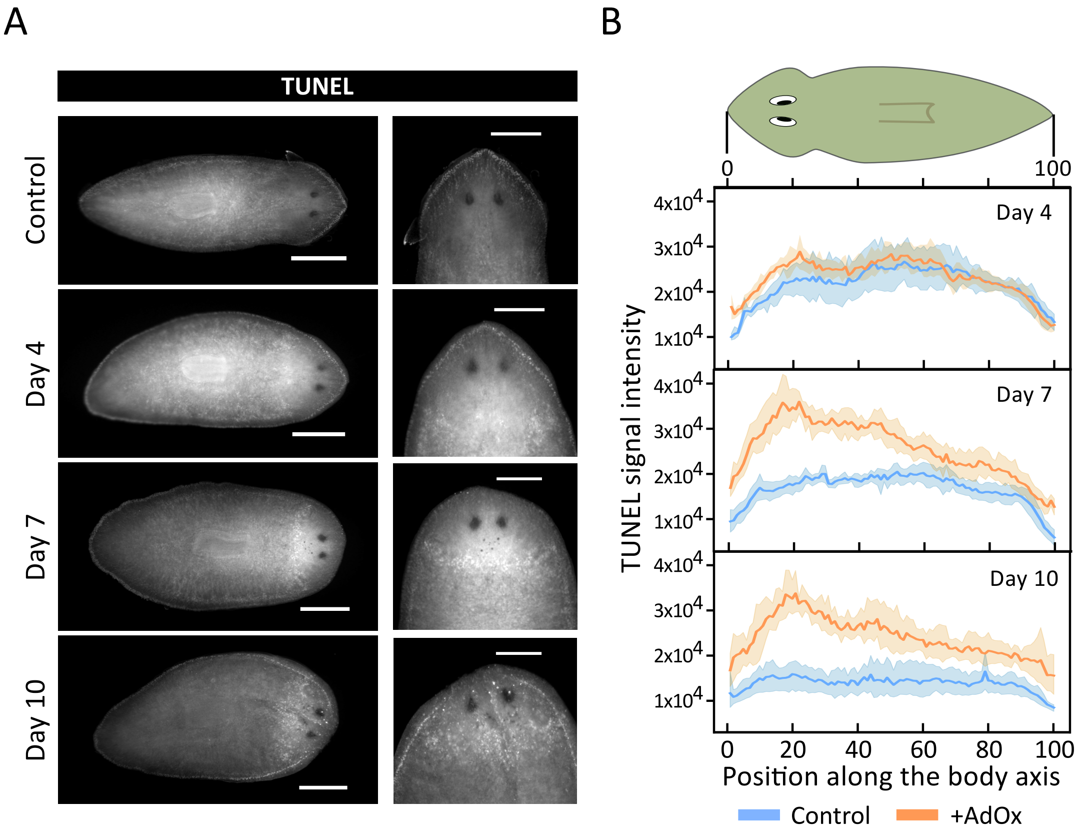

### Fig. S5

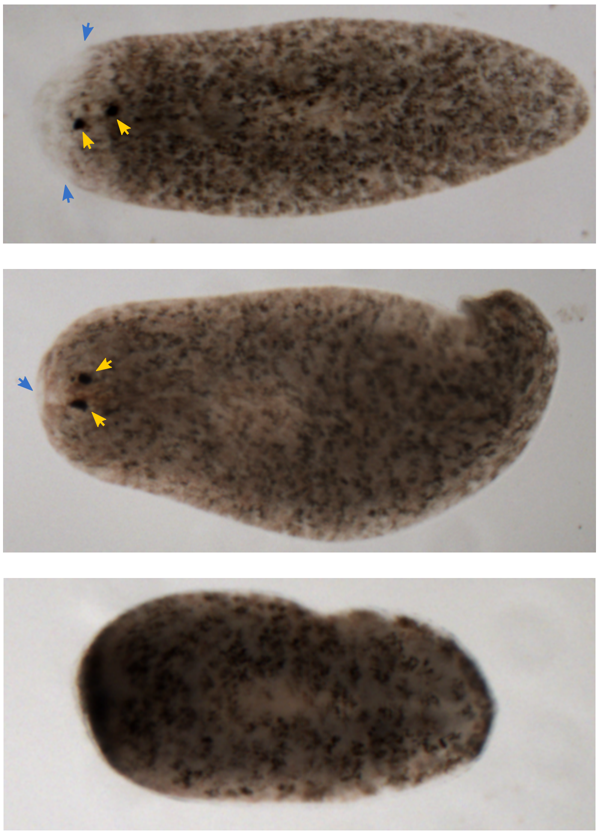

**Figure S5.**

### Fig. S6

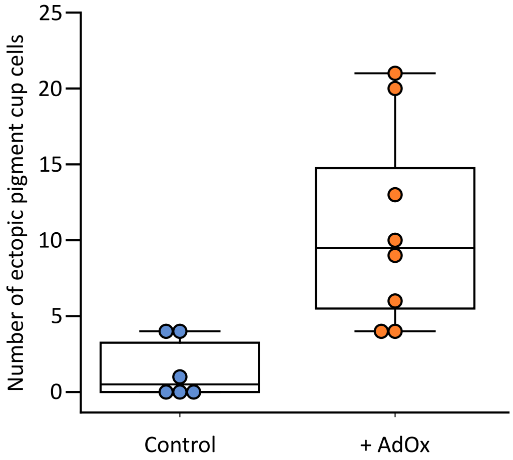

### Fig. S7

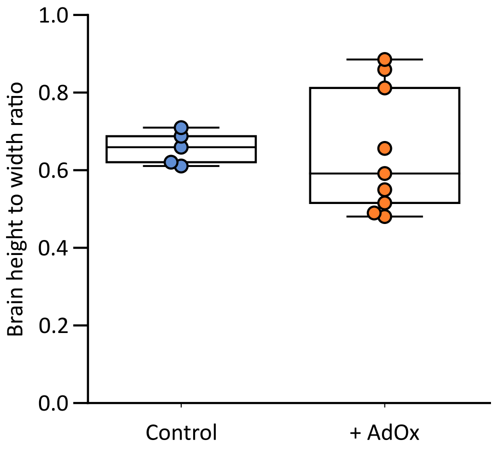


**Figure S7.**

### Fig. S8

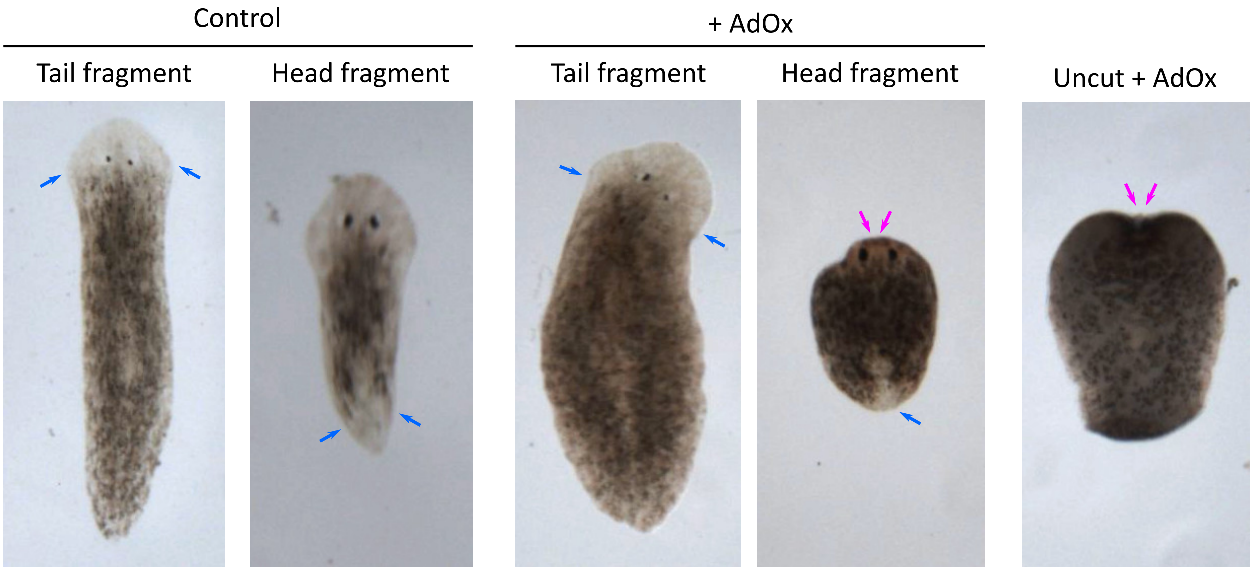

### Fig. S9

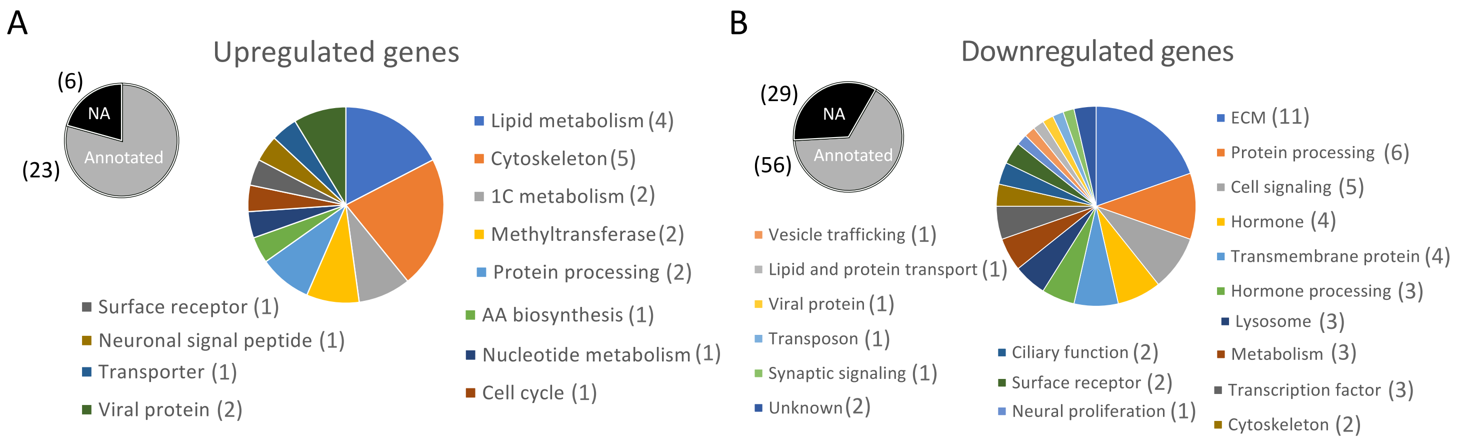

### Fig. S10

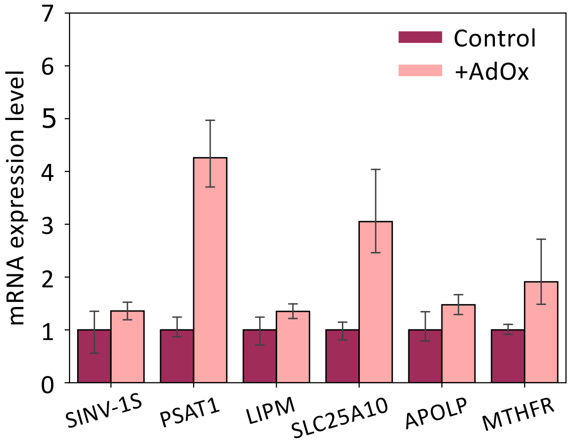

### Fig. S11

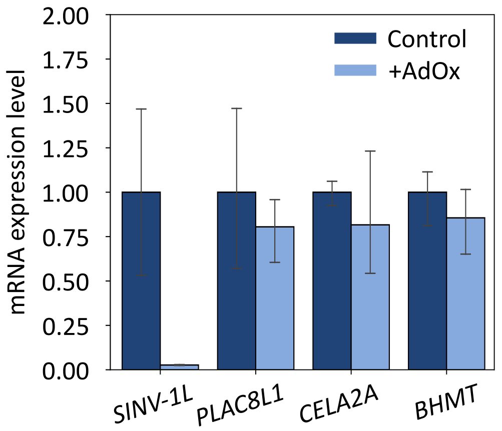

### Fig. S12

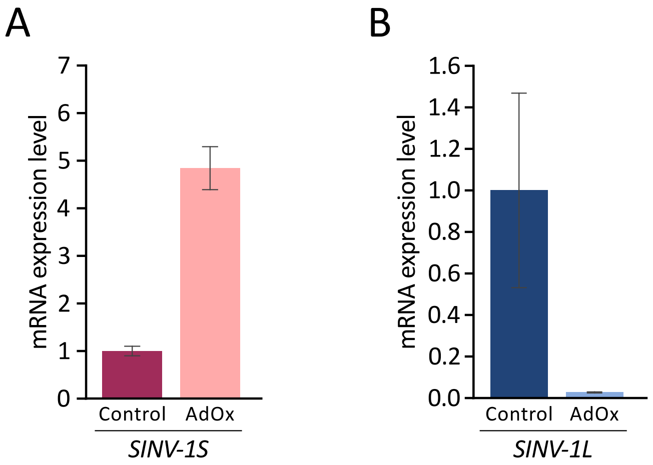
