## Supplemental Data 1 for "S-adenosylhomocysteine hydrolase regulates anterior patterning in *Dugesia japonica*"

**Supplementary figure captions**

**Figure S1. Eye pigment cups of D. japonica are visible after exposure to $H_2O_2$ and light.** Planarians were fixed in Carnoy's fixative and bleached overnight in 6% $H_2O_2$ on a light source. Blue arrows point to the pigment cups. Scale bars – 0.5 mm.

**Figure S2. AdOx does not inhibit cell proliferation in planarians**. A – whole-mount immunohistochemistry showing mitotic cells using anti-H3P antibody in control and AdOx-exposed planarians. Scale bars – 200µm. B – quantification of H3P signal shown in A. Y-axis shows H3P-positive cells per $mm^2$. Ns – Student's t-test, Day 10 p = 0.06.

**Figure S3. AdOx-induced head regression is not due to loss of neoblasts.** A – experimental set-up shown on the left and progression of the head regression in irradiated and AdOx-treated planarians on the right. B – quantitative representation of the head regression and lysis in control and irradiated worms. Number of worms in AdOx and irradiation+AdOx groups – 20, number of worms in irradiated control group – 10.

**Figure S4. Exposure of planarians to SAHH inhibitor AdOx leads to increased anterior apoptosis.** A – TUNEL assay showing apoptotic cells in control and AdOx-treated worms. Scale bars – 200 µm. B – quantification of the TUNEL signal intensity along the body axis. Measurements were done by drawing a line from the head tip to the tail tip to detect the signal intensity. Obtained values were then extrapolated to a 100-point scale for normalization and plotting.

**Figure S5. AdOX-exposed planarians begin tissue regeneration after approximately 3 weeks of drug exposure.** Blue arrows indicate blastema and newly formed tissues. Yellow arrows point at the original eye pigments. Photos were taken on Day 23 of AdOx treatment.

**Figure S6. Number of ectopic cup cells in control and AdOx-treated planarians on Day 32 of the experiment.** Planarians were fixed with Carnoy's fixative and bleached with hydrogen peroxide on a light source overnight to remove non-melanin pigments. Number of ectopic pigment cup cells in AdOx-treated planarians does not include the original pigment cups.

**Figure S7. Brain height to width ratio in control and AdOx-treated planarians on Day 32 of the experiment.** Measurements were done on microscopy images from anti-synapsin immunohistochemistry. AdOx-treated planarian with secondary cephalic connection was not included in this analysis.

**Figure S8. Exposure to SAHH inhibitor AdOx does not impede tissue regeneration.** Images were taken on Day 10 post-amputation. For AdOx treatment, amputated fragments were place in 100 µM AdOx right after the amputation and kept in the same drug solution throughout the experiment. Blue arrows point at the lightly pigmented regenerated tissues. Magenta arrows indicate head regression. Uncut AdOx-treated worm on the right-most image is curled and lies ventral side up.

**Figure S9. Summary of the 119 transcripts identified as differentially expressed in AdOx-adapted planarians.** A – summary of the upregulated genes annotated using planarian database PlanMine 3.0. B – summary of the downregulated genes.

**Figure S10. Independent confirmation of genes identified as significantly upregulated in AdOx-adapted worms.** Quantification of gene expression was done by qPCR using RNA extracts from planarians exposed to AdOx for one month and respective controls. N = 3, error bars – standard error.

**Figure S11. Independent confirmation of genes identified as significantly downregulated in AdOx-adapted planarians.** Gene expression was quantified by qPCR using RNA extracts from planarians exposed to AdOx for one month and from respective controls. N = 3, error bars – standard error.

**Figure S12. Differential effect of AdOx treatment on SINV1 transcript variant expression.** A – qPCR showing upregulation of *SINV-1S* variant in planarians exposed to AdOx for 1 month. B – qPCR of downregulated *SINV-1L* transcript. N = 2, error bars – standard error.
