## Supplementary material for "S-adenosylhomocysteine hydrolase regulates anterior patterning in *Dugesia japonica*": Table S1

| Contig from *Chan et al., 2016* | PlanMine transcript | log2FoldChange | padj | Gene name |
|---|---|---|---|---|
| comp138201_c0_seq1 | dd_Djap_v4_23756_1_1 | -4.276830648 | 9.84E-17 | - |
| comp139701_c0_seq3 | dd_Djap_v4_55278_1_1 | -1.814828736 | 6.05E-05 | - |
| comp125511_c0_seq2 | dd_Djap_v4_27517_1_2 | -1.568222554 | 5.46E-11 | *PLAC8L1* |
| comp125511_c0_seq3 | dd_Djap_v4_27517_1_2 | -1.511265626 | 6.88E-09 | *PLAC8L1* |
| comp133413_c0_seq1 | dd_Djap_v4_21001_1_6 | -1.202347474 | 2.33E-05 | *PXDNL* |
| comp118728_c0_seq1 | dd_Djap_v4_81008_1_7 | -0.997994861 | 6.17E-16 | *PSAP* |
| comp130166_c1_seq1 | dd_Djap_v4_45957_1_1 | -0.742582995 | 7.33E-06 | *CELA2A* |
| comp129491_c1_seq2 | dd_Djap_v4_27179_1_1 | -0.736966678 | 0.001367 | *CELA2A* |
| comp65787_c0_seq1 | dd_Djap_v4_25501_1_1 | -0.732233657 | 0.001242 | *Arrb1* |
| comp129246_c0_seq1 | dd_Djap_v4_58262_1_1 | -0.715548013 | 2.18E-05 | |
| comp122421_c0_seq1 | dd_Djap_v4_50426_1_1 | -0.694934609 | 0.000104 | |
| comp71715_c0_seq1 | dd_Djap_v4_71085_1_1 | -0.687956121 | 1.08E-07 | *FBN2-like* |
| comp127512_c0_seq1 | dd_Djap_v4_72814_1_1 | -0.68277129 | 8.62E-10 | |
| comp115466_c0_seq1 | dd_Djap_v4_27517_1_1 | -0.648133937 | 4.91E-05 | *PLAC8L1* |
| comp116980_c0_seq1 | dd_Djap_v4_45238_1_1 | -0.637061004 | 1.08E-07 | *CTSB* |
| comp125443_c0_seq1 | dd_Djap_v4_72814_1_1 | -0.635320546 | 3.55E-08 | |
| comp137212_c0_seq1 | dd_Djap_v4_30501_1_1 | -0.630368397 | 0.005542 | *Tmem86A-like* |
| comp89256_c0_seq1 | dd_Djap_v4_71085_1_1 | -0.629146864 | 9.09E-09 | |
| comp131951_c0_seq2 | dd_Djap_v4_16855_1_1 | -0.619021401 | 3.55E-05 | *Pam* |
| comp123706_c0_seq1 | dd_Djap_v4_2978_1_1 | -0.603483125 | 0.000839 | *TRAF3* |
| comp147082_c0_seq1 | dd_Djap_v4_58008_1_2 | -0.594637888 | 9.58E-05 | *DUSP10* |
| comp118340_c0_seq1 | dd_Djap_v4_31057_1_1 | -0.590043653 | 6.05E-05 | *NPDC1* |
| comp134449_c0_seq1 | dd_Djap_v4_61342_2_1 | -0.586678793 | 0.002741 | *CMTM4* |
| comp138721_c0_seq2 | dd_Djap_v4_30229_1_1 | -0.554283672 | 0.0005 | *PAM* |
| comp87216_c0_seq2 | dd_Djap_v4_10017_1_1 | -0.545171951 | 0.005655 | *MFGE8* |
| comp104633_c0_seq1 | dd_Djap_v4_55682_1_1 | -0.544880775 | 0.00697 | *CATIP* |
| comp125511_c0_seq1 | dd_Djap_v4_25996_1_1 | -0.544848333 | 0.005016 | *MGC80751* |
| comp132491_c0_seq1 | dd_Djap_v4_24530_2_1 | -0.543757871 | 0.000264 | *PLAC8* |
| comp138721_c0_seq1 | dd_Djap_v4_61786_1_1 | -0.539729396 | 0.001072 | *FBN2* |
| comp88217_c0_seq1 | dd_Djap_v4_34174_1_1 | -0.538892474 | 3.80E-05 | *TSPAN1* |
| comp134449_c1_seq1 | dd_Djap_v4_42359_1_1 | -0.537494486 | 2.69E-05 | *BHMT* |
| comp123693_c0_seq1 | dd_Djap_v4_11226_1_2 | -0.535414017 | 0.005796 | *SNX18* |
| comp143925_c1_seq2 | dd_Djap_v4_3065_1_1 | -0.533240471 | 6.05E-05 | *RSPH1* |
| comp137882_c0_seq1 | dd_Djap_v4_72814_1_1 | -0.526993698 | 9.89E-06 | - |
| comp136024_c0_seq1 | dd_Djap_v4_45490_1_1 | -0.521678034 | 0.002801 | - |
| comp67708_c0_seq1 | dd_Djap_v4_60621_1_1 | -0.518221075 | 0.004546 | - |
| comp131163_c0_seq1 | dd_Djap_v4_47423_1_1 | -0.508719475 | 0.002914 | *P4HB* |
| comp86549_c0_seq1 | dd_Djap_v4_56075_1_1 | -0.503556055 | 0.000338 | *HSPB1* |
| comp130511_c0_seq1 | dd_Djap_v4_56136_1_1 | -0.502711997 | 0.00042 | *HSPA8* |
| comp116100_c0_seq1 | dd_Djap_v4_43021_1_1 | -0.502308535 | 0.000104 | *YBX1* |
| comp67100_c0_seq1 | dd_Djap_v4_81924_1_1 | -0.496137302 | 0.003555 | *7B2-like* |
| comp105199_c0_seq1 | dd_Djap_v4_56037_1_1 | -0.488477807 | 0.000338 | *PCSK2* |
| comp132279_c0_seq1 | dd_Djap_v4_67054_1_1 | -0.48834436 | 0.003951 | *GLIPR1L1* |
| comp137228_c0_seq1 | dd_Djap_v4_68133_1_1 | -0.475427126 | 0.00092 | *CTSB* |
| comp143436_c0_seq5 | dd_Djap_v4_19003_1_1 | -0.458209515 | 0.006518 | *COL1A2-like* |

| | | | | |
|---|---|---|---|---|
| comp134555_c0_seq1 | dd_Djap_v4_9843_1_1 | -0.457124107 | 0.005655 | *COL2A1* |
| comp127851_c0_seq1 | dd_Djap_v4_77732_1_1 | -0.454069859 | 0.000286 | *SEC61B* |
| comp105747_c0_seq1 | dd_Djap_v4_69941_1_1 | -0.452873824 | 0.003118 | *DUSP10* |
| comp141990_c0_seq2 | dd_Djap_v4_98740_1_1 | -0.448986565 | 0.007581 | *SYT1* |
| comp136139_c0_seq1 | dd_Djap_v4_74323_1_1 | -0.444764391 | 0.001442 | *COL5A1-like* |
| comp129214_c0_seq1 | dd_Djap_v4_8987_1_1 | -0.441167125 | 0.001227 | *DNAJA3* |
| comp145471_c0_seq1 | dd_Djap_v4_43021_1_1 | -0.435649766 | 0.001396 | *YBX1* |
| comp115983_c0_seq1 | dd_Djap_v4_32370_1_1 | -0.422307539 | 0.001381 | *NAS-4* |
| comp138721_c0_seq3 | dd_Djap_v4_33592_1_1 | -0.418280248 | 0.003118 | - |
| comp125752_c0_seq1 | dd_Djap_v4_20965_2_1 | -0.416150586 | 0.003785 | *TPPP3* |
| comp102223_c0_seq1 | dd_Djap_v4_26364_1_2 | -0.409353616 | 0.000723 | *HPGDS* |
| comp105626_c0_seq1 | dd_Djap_v4_25300_1_1 | -0.407865486 | 0.000773 | *CNBP* |
| comp84276_c0_seq1 | dd_Djap_v4_52368_2_2 | -0.405176867 | 0.005542 | *DNAI2* |
| comp124681_c0_seq1 | dd_Djap_v4_51680_1_1 | -0.398005104 | 0.002879 | *MAT2A* |
| comp135828_c0_seq1 | dd_Djap_v4_37377_1_1 | -0.39172799 | 0.008883 | *GNL3L* |
| comp135956_c0_seq1 | dd_Djap_v4_65759_1_4 | -0.372190105 | 0.005743 | *HSPA8* |
| comp136104_c0_seq1 | dd_Djap_v4_21261_5_1 | 0.376937406 | 0.006679 | *bt* |
| comp126941_c0_seq1 | dd_Djap_v4_60508_1_1 | 0.40384707 | 0.003134 | *PRSS12* |
| comp147296_c0_seq1 | dd_Djap_v4_31486_3_1 | 0.423142359 | 0.008531 | *MRC1* |
| comp147013_c0_seq1 | dd_Djap_v4_31378_2_1 | 0.469646456 | 0.000595 | *AMN1* |
| comp134863_c0_seq1 | dd_Djap_v4_3862_1_1 | 0.474909011 | 0.000839 | *MYH9* |
| comp129524_c1_seq1 | dd_Djap_v4_73718_1_1 | 0.477669244 | 0.001614 | *PDIA4* |
| comp144449_c0_seq1 | dd_Djap_v4_31486_3_1 | 0.478511342 | 0.001371 | *MRC1* |
| comp130005_c0_seq1 | dd_Djap_v4_14591_1_1 | 0.483792744 | 0.001519 | *ASH1L* |
| comp143779_c0_seq1 | dd_Djap_v4_69987_1_1 | 0.52435229 | 0.001519 | *MTHFR* |
| comp137518_c0_seq1 | dd_Djap_v4_710_2_3 | 0.536209233 | 0.000242 | *HSP90B1* |
| comp85207_c0_seq1 | dd_Djap_v4_47427_1_1 | 0.575618042 | 9.92E-05 | *ASNS* |
| comp138266_c0_seq9 | dd_Djap_v4_64624_1_1 | 0.579173939 | 0.000415 | - |
| comp121199_c0_seq1 | dd_Djap_v4_66233_2_1 | 0.579470138 | 6.05E-05 | *FKBP15* |
| comp145626_c0_seq1 | dd_Djap_v4_31351_1_1 | 0.589410571 | 0.000193 | *APOLTP* |
| comp131320_c0_seq1 | dd_Djap_v4_101130_1_2 | 0.608720955 | 0.009333 | *MACF1* |
| comp125387_c0_seq1 | dd_Djap_v4_80060_1_1 | 0.635611806 | 4.91E-05 | *WBSCR27* |
| comp126941_c0_seq2 | dd_Djap_v4_27300_2_1 | 0.694157714 | 0.009792 | *CG6178* |
| comp67230_c0_seq1 | dd_Djap_v4_11014_2_1 | 0.726076117 | 6.29E-12 | *APOLP* |
| comp105932_c1_seq1 | dd_Djap_v4_35888_2_1 | 0.74614462 | 0.00697 | *TUBB4B* |
| comp131335_c0_seq1 | dd_Djap_v4_36449_1_1 | 0.833721974 | 0.000872 | *SLC25A10* |
| comp124318_c0_seq1 | dd_Djap_v4_2410_1_1 | 1.091128656 | 8.37E-05 | *LIPM* |
| comp128931_c0_seq1 | dd_Djap_v4_18364_1_1 | 1.321779005 | 0.000799 | *PSAT1* |
| comp146649_c0_seq1 | dd_Djap_v4_23756_1_1 | 2.095688675 | 7.24E-16 | - |
| comp139853_c0_seq1 | dd_Djap_v4_13767_1_1 | 2.220776207 | 0.008477 | - |
| comp135977_c0_seq1 | | | | |
| comp135317_c0_seq1 | | | | |
| comp131445_c1_seq1 | | | | |
| comp85890_c0_seq1 | | | | |
| comp143646_c0_seq1 | | | | |
| comp145670_c0_seq2 | | | | |

comp145812_c0_seq4
comp129298_c0_seq1
comp127150_c0_seq1
comp132282_c0_seq3
comp145812_c0_seq3
comp146179_c1_seq1
comp141043_c0_seq1
comp134992_c0_seq1
comp132411_c0_seq1
comp134215_c1_seq1
comp135736_c0_seq1
comp142237_c0_seq1
comp142123_c1_seq1
comp145489_c0_seq2
comp132462_c0_seq1
comp124615_c0_seq1
comp139419_c0_seq3
comp138875_c0_seq1
comp141696_c4_seq2
comp146234_c0_seq1
comp122096_c2_seq1
comp144818_c0_seq1
comp134477_c0_seq2
comp127420_c0_seq1
comp127210_c0_seq1
comp130466_c0_seq2
comp145445_c0_seq1
comp140981_c0_seq1

| Description |
| --- |
| contains putative helicase, protease, RNA-dependent RNA polymerase |
| MARINER TRANSPOSASE |
| PLAC8-like 1 |
| PLAC8-like 1 |
| peroxidasin-like |
| prosaposin |
| chymotrypsin like elastase family member 2A |
| chymotrypsin like elastase family member 2A |
| beta-arrestin-1-like |
| prohormone-4-like, contains LDL receptor-like module |
| perlucin-like protein, C-type lectin superfamily member |
| fibrillin-2-like, contains calcium-bindin EGF domain, EGF/laminin domain |
| contains Mannose 6-phosphate receptor domain |
| PLAC8-like 1 |
| cathepsin B, cystein proteinase 6 containing |
| contains Mannose 6-phosphate receptor domain |
| lysoplasmalogenase-like protein TMEM86A-like, involved in ether lipid metabolism |
| fibrillin-2-like, contains calcium-bindin EGF domain, EGF/laminin domain |
| peptidyl-glycine alpha-amidating monooxygenase A-like |
| TNF receptor-associated factor 3 |
| dual specificity phosphatase 10 |
| neural proliferation differentiation and control protein 1-like |
| CKLF-like MARVEL transmembrane domain-containing protein 4 |
| probable peptidylglycine alpha-hydroxylating monooxygenase |
| milk fat globule-EGF factor 8 protein |
| ciliogenesis associated TTC17 interacting protein |
| tetraspanin |
| placenta-specific 8 |
| fibrillin-2 |
| tetraspanin-1 |
| betaine--homocysteine S-methyltransferase |
| sorting nexin 18 |
| radial spoke head 1 homolog |
| Mannose 6-phosphate receptor domain |
| prohormone-4-like, contains LDL receptor-like module |
| prohormone-4-like, contains LDL receptor-like module |
| prolyl 4-hydroxylase subunit beta |
| heat shock protein beta-1 |
| heat shock protein family A (Hsp70) member 8 |
| Y box protein 1 |
| neuroendocrine protein 7B2-like |
| proprotein convertase subtilisin/kexin type 2 |
| GLI-pathogenesis related 1-like |
| cathepsin B |
| collagen alpha-2(I) chain-like |

collagen, type II, alpha 1
SEC61 translocon beta subunit
dual specificity protein phosphatase 10-like
synaptogamin 11
collagen alpha-1(V) chain-like
DnaJ heat shock protein family (Hsp40) member A3
Y box protein 1
zinc metalloproteinase nas-4-like
C-type lectin mannose-binding isoform-like
tubulin polymerization promoting protein family member 3
hematopoietic prostaglandin D synthase
cellular nucleic acid binding protein
dynein axonemal intermediate chain 2
methionine adenosyltransferase 2A
guanine nucleotide-binding protein-like 3 homolog
heat shock protein family A (Hsp70) member 8
bent, titin
neurotrypsin-like
macrophage mannose receptor 1-like, C-type lectin like
antagonist of mitotic exit network 1 homolog
myosin, heavy chain 9, non-muscle
protein disulfide isomerase family A member 4
macrophage mannose receptor 1-like, C-type lectin like
ASH1 like histone lysine methyltransferase
methylenetetrahydrofolate reductase (NAD(P)H)
Heat shock protein 90kDa beta family member 1
Asparagine synthetase
pseudouridine metabolising bifunctional protein C1861.05-like, PfkB containing protein
Fk506 binding protein 15, 133kDa
Apolipoprotein lipid transfer particle, ApoB
microtubule-actin cross-linking factor 1, isoforms 1/2/3/5-like
Williams-Beuren syndrome chromosomal region 27 protein-like, ubiE/COQ5 methyltransferase
CG6178 gene product from transcript CG6178-RA, fatty acyl-CoA synthethase
apolipophorins-like
tubulin beta 4B class IVb
solute carrier family 25 member 10
lipase, family member M
phosphoserine aminotransferase 1
contains putative helicase, protease, RNA-dependent RNA polymerase
replicase polyprotein

| Organism |
| --- |
| *Solenopsis invicta virus-1* |
| - |
| *Homo sapiens* |
| *Homo sapiens* |
| *Acyrthosiphon pisum* |
| *Homo sapiens* |
| *Homo sapiens* |
| *Homo sapiens* |
| *Crassostrea gigas* |
| *Crassostrea gigas* |
| *Crassostrea gigas* |
| *Lepisosteus oculatus* |
| - |
| *Mus musculus* |
| *Mus musculus* |
| - |
| *Saccoglossus kowalevskii* |
| *Lepisosteus oculatus* |
| *Crassostrea gigas* |
| *Mus musculus* |
| *Mus musculus* |
| *Aplysia californica* |
| *Crassostrea gigas* |
| *Crassostrea gigas* |
| *Mus musculus* |
| *Homo sapiens* |
| *Xenopus laevis* |
| *Homo sapiens* |
| *Homo sapiens* |
| *Tribolium castaneum* |
| *Homo sapiens* |
| *Struthio camelus australis* |
| *Homo sapiens* |
| - |
| *Aplysia californica* |
| *Crassostrea gigas* |
| *Homo sapiens* |
| *Trichinella spiralis* |
| *Homo sapiens* |
| *Mus musculus* |
| *Bombyx mori* |
| *Homo sapiens* |
| *Mus musculus* |
| *Homo sapiens* |
| *Pundamilia nyererei* |

*Heterocephalus glaber*
*Homo sapiens*
*Crassostrea gigas*
*Homo sapiens*
*Megachile rotundata*
*Homo sapiens*
*Mus musculus*
*Vollenhovia emeryi*
*Saccoglossus kowalevskii*
*Homo sapiens*
*Homo sapiens*
*Mus musculus*
*Homo sapiens*
*Homo sapiens*
*Aplysia californica*
*Homo sapiens*
*Drosophila melanogaster*
*Cariama cristata*
*Danio rerio*
*Homo sapiens*
*Anolis carolinensis*
*Homo sapiens*
*Danio rerio*
*Homo sapiens*
*Homo sapiens*
*Homo sapiens*
*Mus musculus*
*Drosophila melanogaster*
*Melopsittacus undulatus*
*Drosophila melanogaster*
*Crassostrea gigas*
*Crassostrea gigas*
*Drosophila melanogaster*
*Crassostrea gigas*
*Homo sapiens*
*Homo sapiens*
*Homo sapiens*
*Mus musculus*
*Solenopsis invicta virus-1*
*Acute bee paralysis virus*
