## Supplementary material for "S-adenosylhomocysteine hydrolase regulates anterior patterning in *Dugesia japonica*": Table S2

| Primer name | Primer sequence |
|---|---|
| Dj_PSAT1-F | 5' TCATCAATACATGCGTACATCATC 3' |
| Dj_PSAT1-R | 5' AATGAACTCAATGTATAATACTCCACC 3' |
| Dj_LIPM-F | 5' GGATTTGATGTATGGCTCAGTAAT 3' |
| Dj_LIPM-R | 5' GATAAAGCATAATTAATTACAGCTGG 3' |
| Dj_SLC25A10-F | 5' CAATGGTCATAAACACAGCTCG 3' |
| Dj_SLC25A10-R | 5' TGCAAAATGATATGAAATTAGCAG 3' |
| Dj_APLPL-F | 5' CGATAATCTATCCATTCAGAGACATA 3' |
| Dj_APLPL-R | 5' GAATCGGTGAAGTATATTTAACAATTAC 3' |
| Dj_MTHFR-1-F | 5' TCACTCCTAAATCTATTCCGTACTG 3' |
| Dj_MTHFR-1-R | 5' CTGGAATTATGCCAATTCAGG 3' |
| Dj_PLAC8L1-F | 5' ACGTAGGAACTGGAGTTCTGGA 3' |
| Dj_PLAC8L1-R | 5' GTAGAACCTGGAACACAAATTGG 3' |
| Dj_CELA2A-F | 5' ATCTGATCTCTGATTTCCTTCCA 3' |
| Dj_CELA2A-R | 5' GGTGTATCTAACATTAACGACTTCG 3' |
| Dj_BHMT-F | 5' GGTTACATGACTCCTGATGCAAC 3' |
| Dj_BHMT-R | 5' ACAGCCTCCAATATATCGAATTC 3' |
